# Humidity as a potential zeitgeber for circadian entrainment of insect systems

**DOI:** 10.1101/2024.09.04.611265

**Authors:** Shyh-Chi Chen, Grace Goodhart, Daniel Eaton, Nathan Catlett, Tabitha Cady, Hannah Tran, Luke E. Lutz, Lyn Wang, Ella Girard, Jaida Savino, Jodi Perry, Libby Hall, Leo Walker, Amena Bidiwala, Emma Tarter, Joshua Tompkin, Nina Greene, Aiden Yang, Joshua B. Benoit

## Abstract

Humidity levels, like light and temperature, fluctuate daily yet are less predictable; however, whether humidity can entrain circadian clocks and synchronize animal behaviors with environmental variations remains unknown. Here, we investigate the circadian humidity entrainment in various insects across multiple orders. Insect species respond to humidity cycles with distinct patterns, some active during wet periods or at the arid-humid transition. When the humidity cue is removed, most species continue to show rhythmic activity associated with the previous arid-humid (AH) cycles. Fruit flies shift their activity accordingly when humidity cycles are altered and remain in the new rhythms under the following free-running conditions (FRC; constant humidity, HH). Moreover, *Drosophila* clock and hygrosensation mutants lack rhythmic activity during (AH) and after humidity entrainment (FRC with HH), indicating that core clock components and hygrosensors are essential for circadian entrainment. Our findings provide strong evidence that humidity is likely to serve as a potential zeitgeber for circadian entrainment in most, but not all, insect systems and will likely have broad applicability and importance across animal systems. While light and temperature act as the primary zeitgebers, understanding the mechanisms of humidity entrainment will help us better interpret the behavioral patterns of terrestrial animals, particularly those susceptible to dehydration.

**One Sentence Summary:** Humidity entrainment of the circadian clock synchronizes insect activity to environmental changes.

## Introduction

Animals experience daily fluctuations in their surroundings, mainly driven by changes in light levels and temperature. These daily cycles of light and temperature are the main environmental cues (zeitgebers) that entrain the circadian clock and synchronize organismal biological functions ^1,2^. Circadian disruption and misalignment often lead to harmful effects on organisms ^3,4^. Studies on insects have shown that, besides light and temperature, multiple biotic and abiotic factors play important roles in the synchronization of circadian rhythms ^5–8^. One specific abiotic factor that has not been broadly examined is whether humidity can serve as a potential zeitgeber in animals. Of interest is that humidity has distinct daily cycles, but does show high variability based on precipitation, and terrestrial animals often show adaptive responses to humidity changes to avoid desiccation ^9,10^. For terrestrial arthropods, humidity affects the spatiotemporal dynamics of behavior and physiology. Exposure to arid conditions alters the general activity and feeding behaviors of various pest species ^11,12^. Of particular interest is that several factors that underlie humidity sensing have been recently identified in insects ^13–21^. There is growing evidence on how animals respond to changes in humidity, including a few studies focused on the interaction between hygrosensation and diel activity in insects and other animals. For example, a previous study investigated whether humidity could act as a zeitgeber for *Drosophila* eclosion but found that *Drosophila* could not be entrained to humidity cycles and eclosed arrhythmically ^22^. Interestingly, a study in plants shows that humidity oscillations can entrain the circadian clock in *Arabidopsis* in constant light (LL) and light-dark cycles (LD), with higher expression amplitude of clock genes in the latter condition ^23^. However, whether humidity can entrain the circadian activity in animal systems remains unclear.

This study aims to examine how daily humidity cycles influence circadian synchronization for insects. First, entrainment under arid-humid (AH) cycles followed by free-running conditions (FRC; constant humidity, HH) was examined for insect species across three orders. These studies were followed by targeted *Drosophila* studies to establish the relationship between cycling humidity and activity patterns and examine whether mutant lines with defects in humidity detection and clock processes can behaviorally synchronize to humidity cycles. These studies provide evidence of a novel humidity-entrainable clock in insects that requires humidity sensing and clock genes that influence circadian biology of insects. The identification of humidity as a zeitgeber will help better predict how insect behavior and distribution may be altered by environmental conditions.

## Results

### Humidity cycles can entrain insect locomotor activities

Previous studies have shown that temperature cycles can synchronize behavioral activity and clock gene expression under both constant light (LL) and constant darkness (DD) ^24,25^. Therefore, we ask whether the activities of various arthropods can be synchronized to humidity cycles and whether humidity can entrain circadian clocks under LL and DD. To establish if humidity cycling can entrain circadian clocks, the activity pattern of various insect species under arid (A, 40-50%RH)-humid (H, > 85%RH) cycles (12h:12h A:H) was examined with stable temperature (25°C), and air pressure (780 mm Hg) under LL (350 lux) or DD conditions (Fig. 1-2; Supplementary Fig. 1). Under these conditions, there were only daily variations in humidity with temperature, light, and barometric pressure showing no differences. In the present study, all insects examined robustly respond to environmental humidity changes (AHLL) with two distinct behavioral patterns and varying activity levels (Fig. 2A-C; Table 1). Kissing bugs, spider beetles, and two mosquito species (daytime-active *Aedes aegypti* and nighttime-active *Culex pipiens*) showed higher activity levels in the dry phase than in the humid phase (Fig. 2C). Importantly, these behavioral shifts were rather immediate, within 30-60 minutes of the shifts from wet to dry conditions. These rapid shifts suggest that activity differences are due to dry conditions rather than dehydration, which has been observed in multiple systems ^11,26,27^. *Drosophila* exhibited a behavioral pattern with an activity peak at the arid-to-humid transition (Fig. 2B, near ZT 12).

**Fig. 1.**
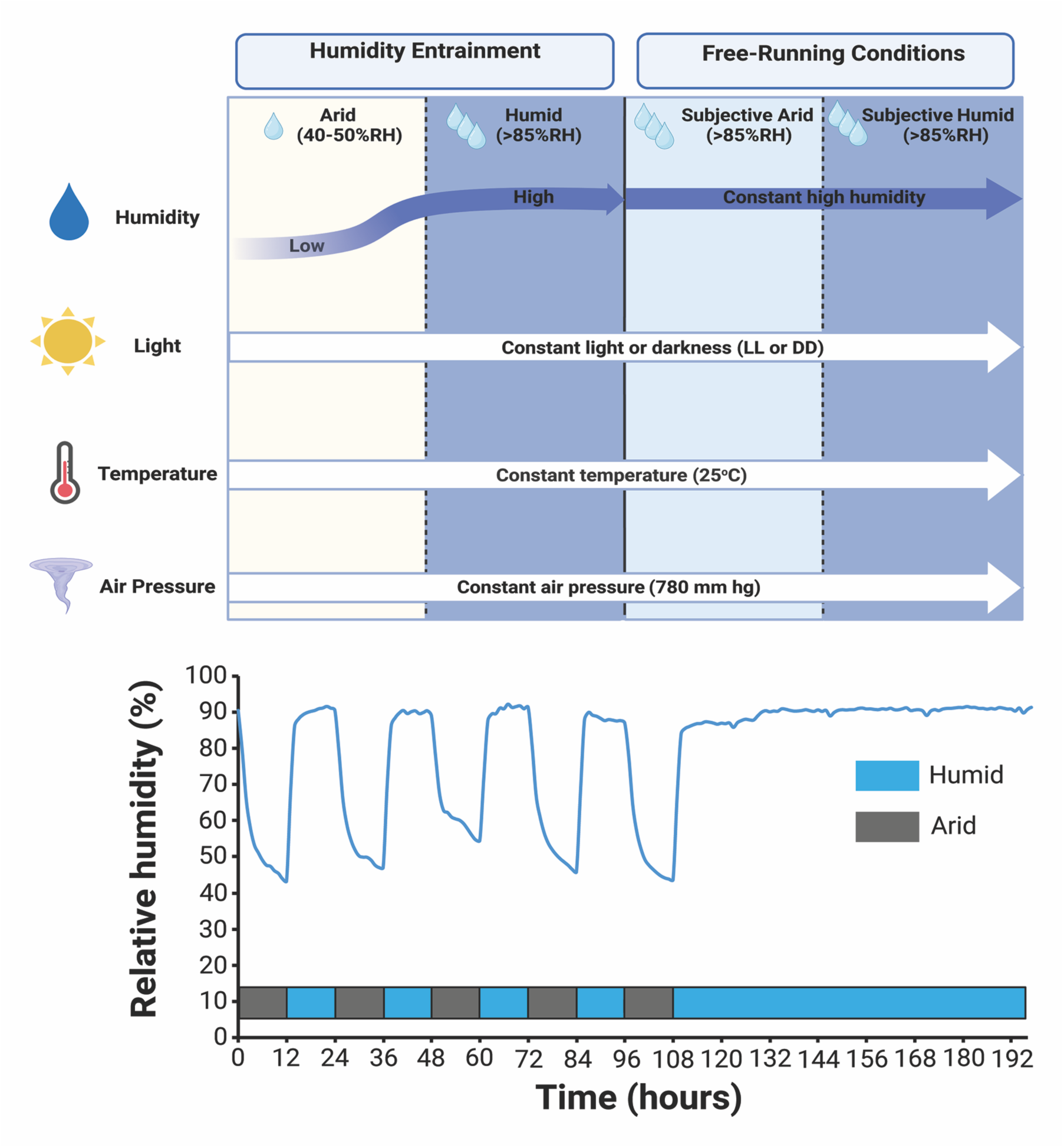
Humidity entrainment of insect circadian clocks. The schematic shows how insects are entrained in the humidity cycles (12h A:12h H) for 5 days and then in the free-running conditions with constant humid (HH) conditions. Light, temperature, and air pressure remain constant during experiments.

**Fig. 2.**
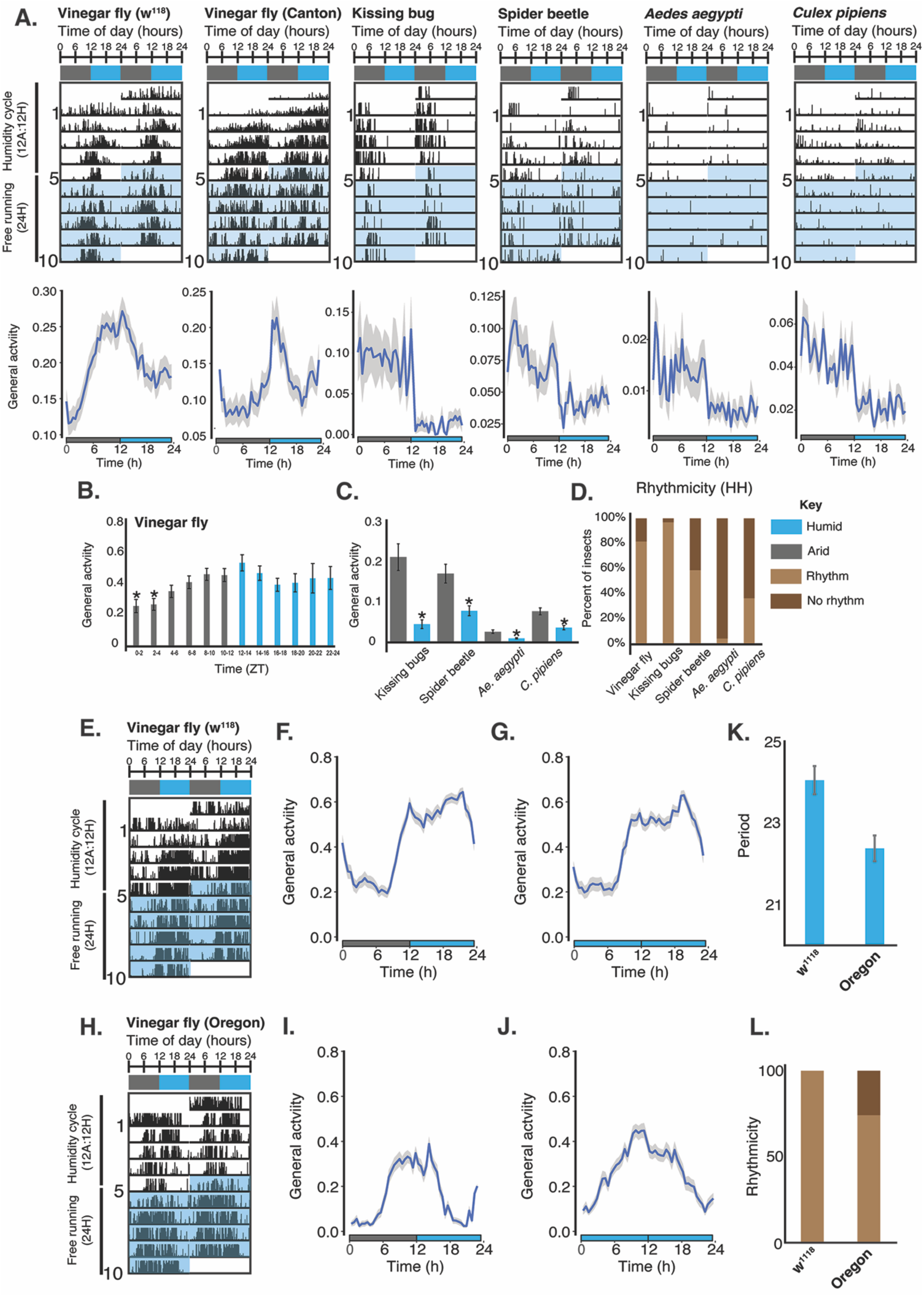
The humidity entrainment of circadian clocks of various insect species. (A) Top: Double-plotted actograms of insects in the humidity cycles and free-running conditions. Bottom: General activity patterns of insects in the last two days of the humidity cycles. Numbers of animals examined (n): vinegar flies, *Drosophila melanogaster* (*w*^1118^, ♂=36 and Canton S, ♂=35), kissing bugs, *Rhodnius prolix*us (♀) =33, spider beetles, *Mezium affin*e (sex unknown) =56, yellow fever mosquitoes, *Ae. aegypt*i (♀) =50, northern house mosquitoes, *C. pipiens* (♀) =60. (B) Twenty-four-hour activity of *Drosophila w*^1118^ during the humidity entrainment (n=115). One-way ANOVA and post hoc Tukey HSD test were used to compare with ZT12–14. ∗ denotes p < 0.05. (C) Comparison of activities of multiple insects in the dry and humid phases during the humidity entrainment. n for each species: kissing bugs=19, spider beetles=71, *Ae. aegypt*i=50, *C. pipiens*=59. ∗ denotes p < 0.05 in Student’s t-test. (D) Rhythmicity of various insects in the free-running conditions. Numbers above the bar graph: n for each species. (E-G) *w*^1118^: representative actogram (E), general activity patterns of humidity cycles (F, n=44) and FRC (G, n=47). (H-J) Oregon-R-C: representative actogram (H), general activity patterns of humidity cycles (I, n=26) and FRC (J, n=23). (K,L) Period (K) and Rhythmicity (L) of *w*^1118^ (n=47) and Oregon-R-C (n=23) in the FRC. (A-D) Flies were raised for their entire life under LD and moved to LL for the humidity assay. (E-L) Flies were raised for their entire life under LL and moved to DD for the humidity assay.

**Table 1.** Locomotor activity rhythms of various insect species in AHLL and HHLL.

| Species | Humidity Cycles (AHLL) |  |  | Free-Running Conditions (HHLL) |  |  |
| --- | --- | --- | --- | --- | --- | --- |
|  | Rhythmicity<br>(%, n<sup>R</sup>/n) | Power | Period<br>(h) | Rhythmicity<br>(%, n<sup>R</sup>/n) | Power | Period<br>(h) |
| *D. melanogaster*<br>(*w*<sup>1118</sup>, ♂) | 83.3<br>(30/36) | 167.7 ±<br>19.0 | 23.7 ±<br>0.2 | 62.9<br>(22/35) | 187.4 ±<br>29.2 | 24.8 ±<br>0.6 |
| *D. melanogaster*<br>(Canton S, ♂) | 85.7<br>(30/35) | 192.3 ±<br>19.2 | 23.5 ±<br>0.1 | 74.3<br>(26/35) | 131.6 ±<br>20.8 | 24.5 ±<br>0.6 |
| *R. prolixus* (♀) | 85<br>(17/20) | 571.3 ±<br>85.4 | 24.0 ±<br>0.4 | 96.97<br>(32/33) | 462.18 ±<br>42.28 | 25.12 ±<br>0.72 |
| *M. affine*<br>(sex unknown) | 76.8<br>(43/56) | 350.9 ±<br>40.7 | 24.5 ±<br>0.3 | 58.97<br>(23/39) | 279.19 ±<br>41.73 | 25 ±<br>0.78 |
| *Ae. aegypti* (♀) | 12<br>(6/50) | 221.7 ±<br>96.0 | 24.1 ±<br>1.4 | 4.65<br>(2/43) | 392.74 ±<br>73.63 | 30.8 ±<br>0.47 |
| *C. pipiens* (♀) | 34.0<br>(18/53) | 392.8 ±<br>50.3 | 24.8 ±<br>0.4 | 36.67<br>(22/60) | 348.87 ±<br>57.69 | 25.16 ±<br>0.81 |

n, number of flies analyzed; n<sup>R</sup>, rhythmic flies.

To determine whether humidity can serve as a zeitgeber to entrain the circadian clock, we examined insect activity under constant humidity (HHLL) following entrainment under humidity cycles (AHLL). Except for mosquitoes, all other insects showed high levels of rhythmic behavior when shifted from AHLL to constant humidity conditions (HHLL, Fig. 2D). We observed that flies raised for their entire life in LL under constant temperature also showed rhythmic behaviors in the humidity cycles (AH) and free-running (HH) in constant dark conditions (DD) (Fig. 2E-G: *w*^1118^; Fig.2H-J: Oregon-R-C; Fig. 2K-L, and Table 2: period and rhythmicity in the free-running conditions, FRC, HH). Advancing the humidity phase under LL by 6 hours led to a behavioral shift to match the new cycle within 3 days, indicating entrainment to changes in this cue (Fig. 3). These results indicate that terrestrial insects respond to daily humidity fluctuations in a manner that effectively synchronizes their behavior with these fluctuations.

**Fig. 3.**
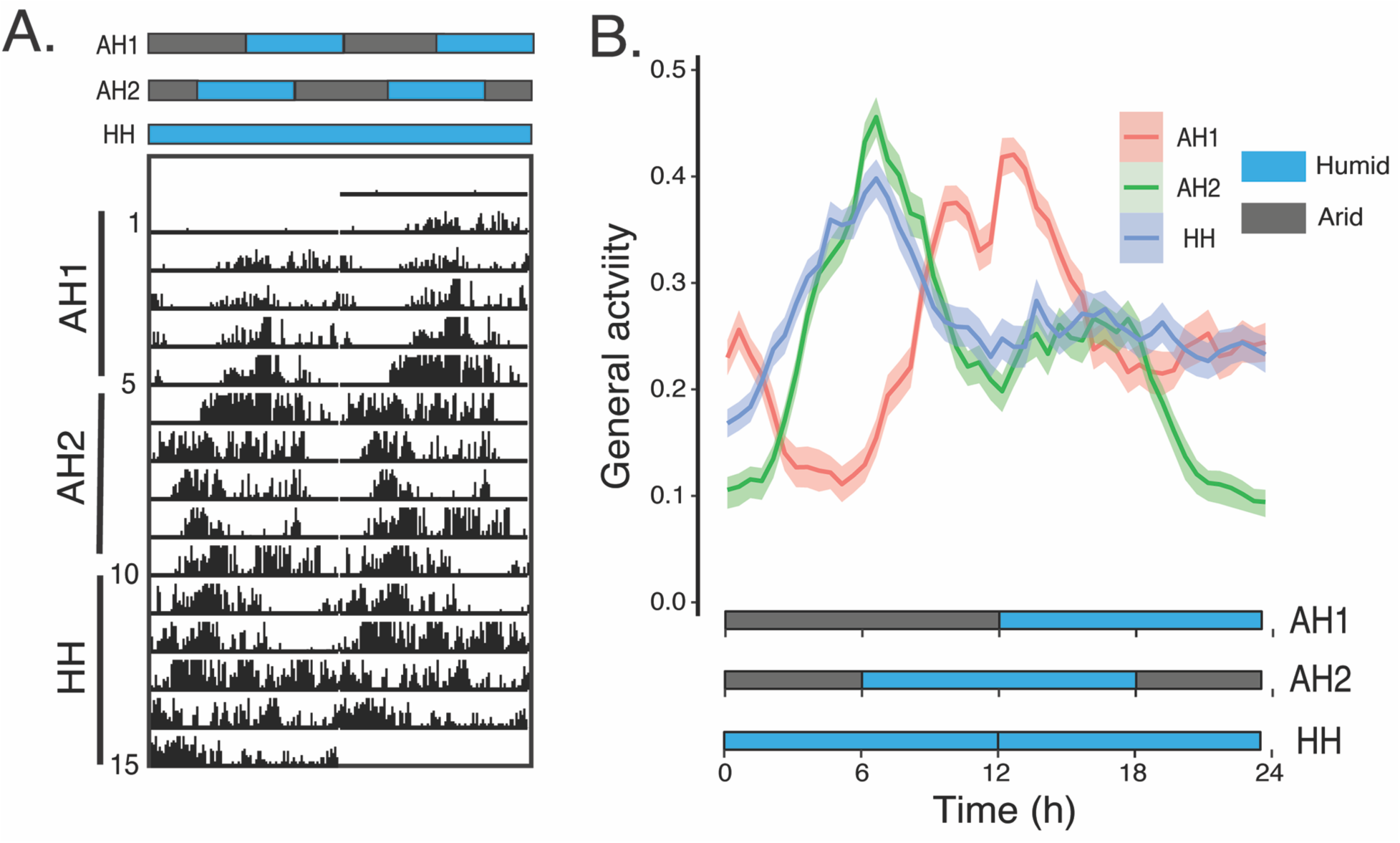
The shift in fly activity in response to the change in the humidity regime. (A) A representative actogram of a *Drosophila w*^1118^. (B) The curves represent the average activity patterns of *w*^1118^ flies (♂, n=62) in two humidity regimes (AH1 and 6h phase-advanced AH2) and the following free-running condition (HH). All experiments are under LL. Bar colors represent different phases: arid (Dark gray) and humid (blue).

**Table 2.** Locomotor activity rhythms of various insect species in AHDD and HHDD.

| Species | Humidity Cycles (AHDD) |  |  | Free-Running Conditions (HHDD) |  |  |
| --- | --- | --- | --- | --- | --- | --- |
|  | Rhythmicity<br>(%, n<sup>R</sup>/n) | Power | Period<br>(h) | Rhythmicity<br>(%, n<sup>R</sup>/n) | Power | Period<br>(h) |
| *D. melanogaster*<br>(*w*<sup>1118</sup>, ♂) | 93.6<br>(44/47) | 282.1 ±<br>22.3 | 24.0 ±<br>0.2 | 93.6<br>(44/47) | 292.9 ±<br>24.1 | 24.1 ±<br>0.4 |
| *D. melanogaster*<br>(Oregon-R-C, ♂) | 77.4<br>(24/31) | 211.9 ±<br>25.9 | 23.8 ±<br>0.2 | 67.7<br>(21/31) | 144.4 ±<br>29.1 | 22.8 ±<br>0.2 |

n, number of flies analyzed; n<sup>R</sup>, rhythmic flies.

### Humidity-associated activities are under circadian control

After establishing a distinct humidity-cycle response and entrainment, we aimed to determine whether insect humidity-associated activities are under circadian regulation. We examined the activities of *Drosophila* mutants for key clock genes (*per*^01^*, tim*^01^*, cyc*^01^, and *Clk^Jrk^*) and the main circadian neuropeptide (*pdf*^01^) under AH and HH conditions. Although these mutants responded to the humidity changes under AH to some extent (Fig. 4A), all lines exhibited weaker rhythmicity with more arrhythmic flies in the FRC compared to *w*^1118^ (Fig. 4B; Table 3). These results suggest that functional circadian clock molecular machinery is essential for humidity entrainment.

**Fig. 4.**
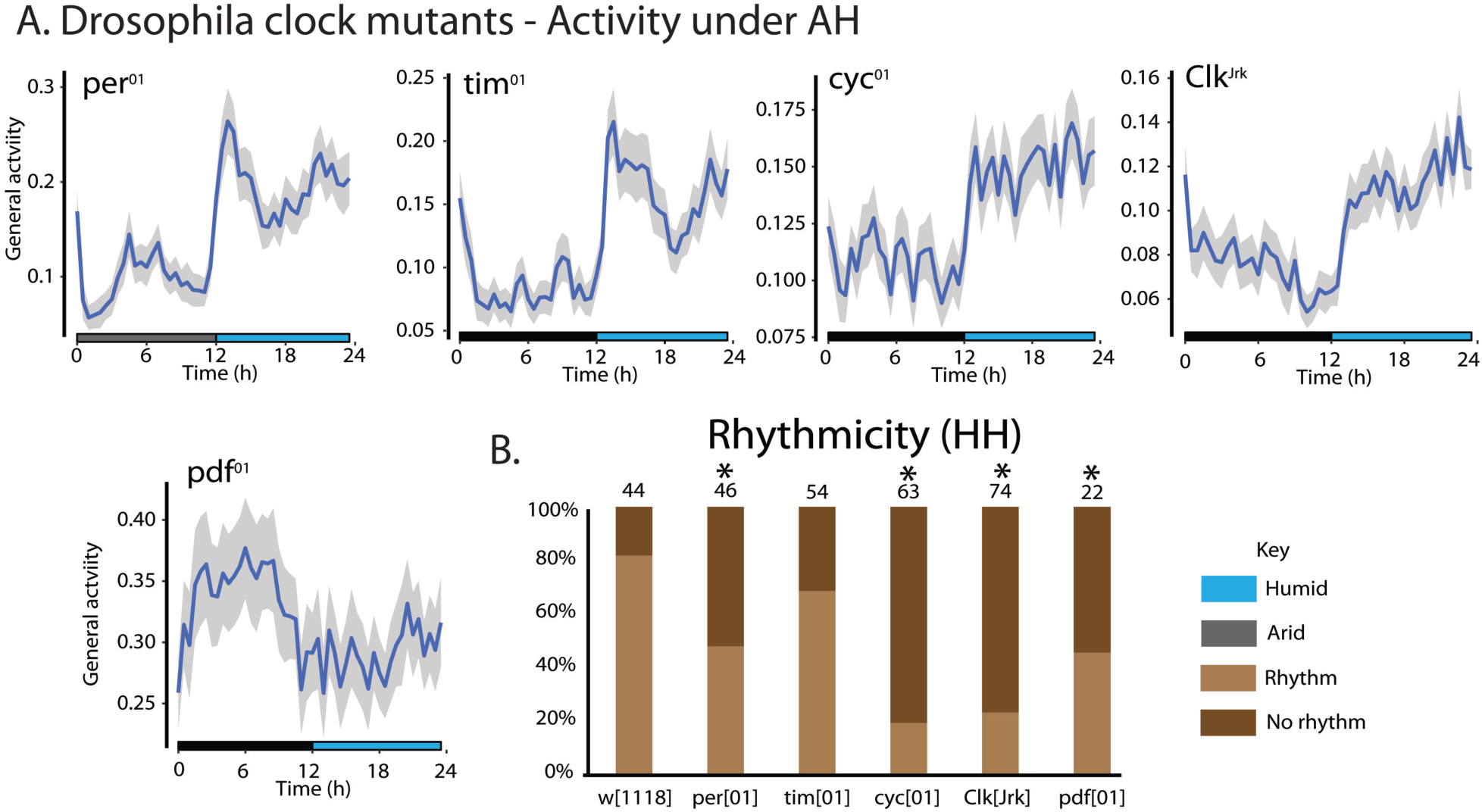
The rhythmic humidity-associated behaviors are under circadian control. (A) General activity patterns of *Drosophila melanogaster* clock and *pdf*^01^ mutants (♂) in the last two days of the humidity cycles. n: *per*^01^=49, *tim*^01^=58, *cyc*^01^=62, *Clk^Jrk^*=75, *pdf*^01^=29. (B) Rhythmicity of clock and *pdf*^01^ mutants in the free-running conditions. All experiments are under LL. Numbers above the bar graph: n in each species. ∗ denotes significance (p < 0.05) in the Fisher exact test followed by the post-hoc Bonferroni test.

**Table 3.** Locomotor activity rhythms of various *Drosophila* clock mutants in HHLL.

| Flies | n | Rhythmicity<br>(%, n<sup>R</sup>/n) |
| --- | --- | --- |
| *w*<sup>1118</sup> (♂) | 44 | 81.82 (36/44) |
| *per*<sup>01</sup> (♂) | 46 | 47.83 (22/46) |
| *tim*<sup>01</sup> (♂) | 54 | 68.52 (37/54) |
| *cyc*<sup>01</sup> (♂) | 63 | 19.05 (12/63) |
| *Clk*<sup>Jrk</sup> (♂) | 74 | 22.97 (17/74) |
| *pdf*<sup>01</sup> (♂) | 22 | 45.45 (10/22) |

n, number of flies analyzed; n<sup>R</sup>, rhythmic flies.

### Humidity sensing-associated molecules are essential for the circadian humidity entrainment

Hygrosensation requires specific sensors ^13,14,16^, which we targeted to determine if these are critical to humidity entrainment. *Drosophila* hygrosensation mutants differentially responded to humidity cycles (Fig. 5A) and most showed lower rhythmicity in HH after entrainment under AH (Fig. 5B). Among them, the rhythmicity of *Ir40a* and *Obp59a* mutants, along with *Ir76b,* which is related to low salt and amino acid, sour sensing ^28–30^, was significantly lower than that of *w*^1118^ flies (Fig. 5B; Table 4). These results reveal that hygrosensation is essential for humidity entrainment of the circadian clock, and insects may use different repertoires of hygrosensing molecules for behavioral synchronization to daily humidity changes. Lack of specific hygrosensing molecules such as Ir40a and Obp59a may impede the input of the humidity signal to the clock, thereby dampening internal molecular oscillations and ultimately leading to behavioral synchronization failure.

**Fig. 5.**
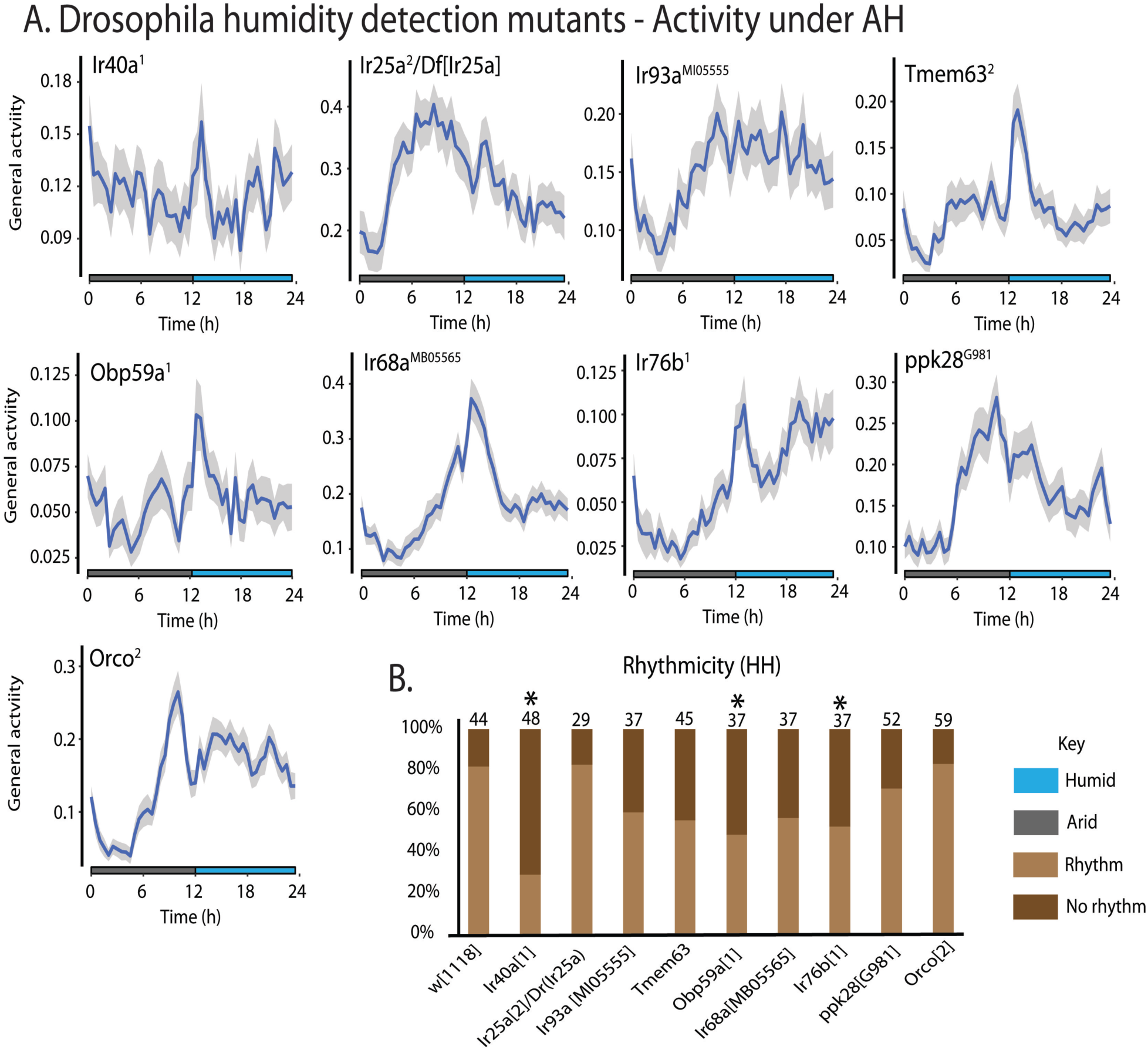
Hygrosensation is required for the humidity entrainment of circadian clock. (A) General activity patterns of *Drosophila* hygrosensation mutants (♂) in the last two days of the humidity cycles. n: *Ir40a*^1^= 55, *Ir25a*^2^*/Df(Ir25a)*=30, *Ir93a^MI05555^*=57, *Tmem63*^2^=45, *Obp59a*^1^=46, *Ir68a^MB05565^*=48, *Ir76b*^1^=60, *ppk28^G981^*=52, *Orco*^2^=66. (B) Rhythmicity of hygrosensation mutants in the free-running conditions. All experiments are under LL. Numbers above the bar graph: n in each species. ∗ denotes significance (p < 0.05) in the Fisher exact test followed by the post-hoc Bonferroni test.

**Table 4.** Locomotor activity rhythms of various *Drosophila* hygrosensation mutants in HHLL.

| Flies | n | Rhythmicity<br>(%, n<sup>R</sup>/n) |
| --- | --- | --- |
| *w<sup>1118</sup>* (♂) | 44 | 81.82 (36/44) |
| *Ir40a<sup>1</sup>* (♂) | 48 | 29.17 (14/48) |
| *Ir25a<sup>01</sup>/Df(Ir25a)* (♂) | 29 | 82.76 (24/29) |
| *Ir93a<sup>MI0555</sup>* (♂) | 37 | 59.46 (22/37) |
| *Tmem63<sup>2</sup>* (♂) | 45 | 55.56 (25/45) |
| *Obp59a<sup>1</sup>* (♂) | 37 | 48.65 (18/37) |
| *Ir68a<sup>MB05565</sup>* (♂) | 37 | 56.76 (21/37) |
| *Ir76b<sup>1</sup>* (♂) | 59 | 52.54 (31/59) |
| *ppk28<sup>G981</sup>* (♂) | 52 | 71.15 (37/52) |
| *Orco<sup>2</sup>* (♂) | 59 | 83.05 (49/59) |

n, number of flies analyzed; n<sup>R</sup>, rhythmic flies

### Hygrosensation and circadian clocks are critical for humidity entrainment of sleep processes

Sleep, similar to locomotor activity, in light:dark cycles is driven by circadian clocks^31^. To understand how circadian humidity entrainment influences sleep, we analyzed the *Drosophila* sleep under AH and HH conditions. Importantly, sleep patterns are not a direct inverse of activity, but are related, e.g. high periods of activity will have reduced sleep levels. The sleep of *w*^1118^ flies showed a peak at the beginning of the arid phase near ZT0 and a trough appeared near the arid-to-humid transition (ZT12) (Supplementary Fig. 2A). On the other hand, the clock and hygrosensor mutants (*per*^01^: Supplementary Fig. 2, B and E; *Ir40a*^1^: Supplementary Fig. 2, C and F) showed different sleep patterns in AH (Supplementary Fig. 2, B and C) and loss of sleep rhythm in the HH compared to *w*^1118^ (Supplementary Fig. 2, D-F). These results suggest that *Drosophila* sleep is driven by the humidity-entrainable circadian clock, which was expected based on our observations of activity.

**Fig. 6.**
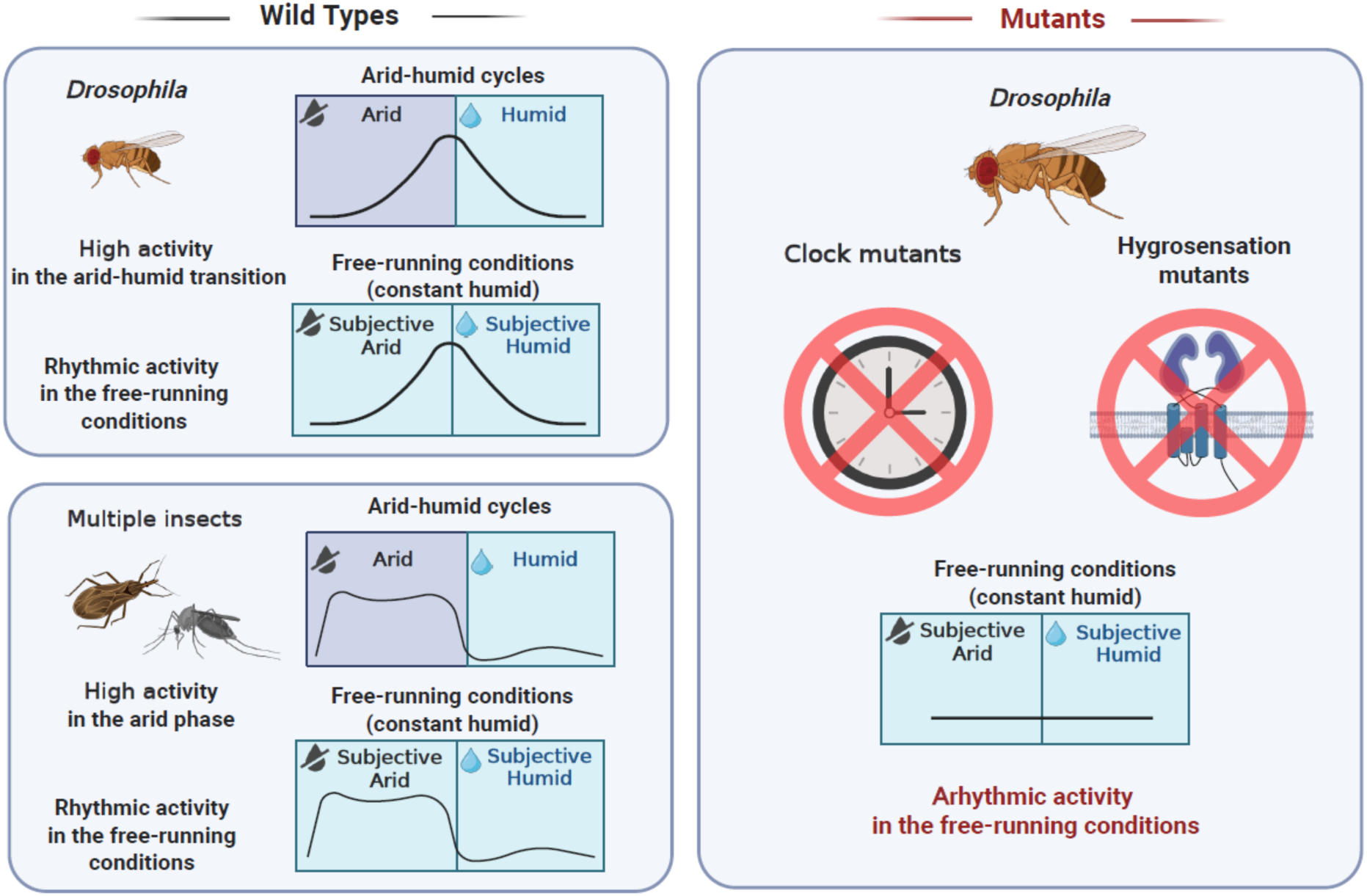
Activity patterns of insects in the humidity cycles and free-running conditions. The schematic shows insect activity after the 5 days of entrainment to humidity cycles (12h A:12h H), followed by free-running conditions with constant humidity. Light, temperature, and air pressure remain constant during experiments. Activity patterns of various insect wild types (left top, *Drosophila*; left bottom, other insects) and *Drosophila* mutants (right).

## Discussion

This study reveals that humidity can serve as a zeitgeber, synchronizing insect behaviors with environmental changes. Outside this study, direct observation of humidity as a potential zeitgeber has been reported in only a single plant system ^23^. Previous literature has suggested that humidity cycles may more closely match insect activity than light:dark or temperature cycles ^32^. In our studies, all insect species respond robustly to humidity cycles, and locomotor activity of some species remains rhythmic in the FRC. *Drosophila* clock and hygrosensation mutants exhibit abnormal phenotypes during humidity entrainment and low rhythmicity under free-running conditions, suggesting that humidity entrainment is under circadian control and requires hygrosensation-associated receptors or molecules (Fig. 5). The humidity-sensing molecules are primarily expressed in antenna ^13–16,19^. Among them, we found that Ir40a and Obp59a may play an important role in the sensory input to this humidity-entrainable clock (Fig. 5). Since both Ir40a and Obp59a are selectively expressed in the sacculus of the antenna ^14,15,15,16^ and the cells expressing Ir40a and Obp59a are adjacent to each other ^16^, the external signal input for circadian entrainment is likely through the neural circuit composed of hygrosensitive neurons in the second chamber of the sacculus, small glomerulus at posterior antennal lob (PAL), and projection neurons relaying the information to the higher brain ^15^.

Circadian clock governs a broad aspect of animals’ lives through the synchronization of organisms to environmental changes. While light and temperature are generally acknowledged as the most potent zeitgebers for circadian rhythms, humidity remains poorly understood in this context and may also play a crucial role in the rhythmic patterns of biological processes, particularly in terrestrial animals. On a warm, dry, sunny day, terrestrial animals gradually experience water loss, which threatens their survival if essential adaptations do not occur, such as rehydration from food sources or reduction of water loss ^11,26,33^. An endogenous humidity-entrainable clock helps animals more accurately predict daily humidity fluctuations. Unlike the well-studied sensory inputs for light and temperature-entrainment and the interaction between light- and temperature-entrained clocks, the humidity-input pathway for circadian rhythms in insects, along with other animals, remains unknown. Furthermore, understanding how humidity interacts with light and temperature could provide new insights into circadian regulation of animal behavior and physiology in response to multiple environmental cues in nature. Most likely, the humidity entrainment represents a much weaker cue compared to light cues, which could increase normal patterns of fruit fly activity, like the observations of light and temperature dynamics ^34^. Daily fluctuations in abiotic environmental factors can be highly correlated (in phase) or out of phase, leading to various impacts on biological processes ^35–38^. Specific experiments will be required to establish how humidity can impact the two most potent clock-resetting signals, temperature and light cycles, which can be influenced by shifting humidity, where it can act in tandem to coordinate activity or act in conflict to impair typical entrained behaviors.

The discrepancy between our finding of behavioral circadian entrainment to humidity and the earlier report that *Drosophila* could not be entrained by humidity cycles ^22^ likely reflects strong stage- and trait-specific differences in circadian sensitivity. The previous study examined adult eclosion timing ^22^, a developmental output that is primarily organized during the pupal stage and is known to be dominated by photoperiodic and thermal cues ^39–42^, potentially limiting its responsiveness to hygric signals. In contrast, our study quantified active adult behavior; a labile adult trait may be under stronger selective pressure to track daily humidity fluctuations to maintain water balance. Sensory pathways, central clock, and peripheral oscillators can differ markedly across life stages ^43–45^, and hygric sensitivity may be heightened in adults relative to immobile or encapsulated developmental stages. Thus, while humidity cycles may be insufficient to entrain developmental timing mechanisms such as eclosion, they can nonetheless serve as effective zeitgebers for adult behavioral rhythms, underscoring the importance of considering both life stage and circadian output when evaluating environmental entrainment.

The lack of humidity entrainment in mosquitoes is surprising due to the close relationship to fruit flies ^20,46,47^. Humidity is unlikely to serve as a zeitgeber in mosquitoes as it functions as a host cue ^20,47,48^. Elevated humidity gradients are tightly associated with vertebrate hosts through respiration and perspiration and are used by mosquitoes for orientation toward blood meal sources along with other cues ^20,49^. As humidity signals are strongly contingent on host presence, they lack the predictability required of a reliable zeitgeber for mosquitoes. As such, mosquitoes appear to interpret humidity as an immediate sensory cue that modulates host-seeking on short timescales, rather than as a temporal signal capable of synchronizing circadian rhythms in relation to daily cycles. This lack of entrainment suggests that not all insects use humidity as a zeitgeber, and that this input system needs examination across specific insect taxa.

The humidity-entrainable oscillator likely exists widely among various terrestrial arthropods. Synchronizing behaviors to environmental humidity changes is essential for animal fitness, particularly for avoiding dehydration ^11,50,51^. This study paves the way to explore the relationship between daily humidity fluctuations and the activities of many pests. Future studies that explore the specific molecular and neuronal aspects essential for these humidity cycles to act as a zeitgeber are required to elucidate the mechanisms of this process. For example, whether the previously reported neuronal network for sensory integration of temperature and humidity in the brain is employed for humidity entrainment ^15^. Our observation that clock and humidity detection genes are critical to this entrainment does provide evidence that peripheral humidity detection and circadian clock genes are critical to humidity-entraining insect systems. Understanding the underlying mechanism will help us not only predict the time-of-day behavioral patterns of numerous harmful agricultural and public health pests but also develop effective strategies to reduce vector-host interactions for disease control.

## Methods

### Animals and husbandry

In most experiments (AHLL/HHLL) of this study, animals were reared in the insectary under 15h:9h Light:Dark cycles, 25°C and 70-80% relative humidity (RH). For experiments in AHDD/HHDD conditions, flies were reared under constant light conditions (LL) for their entire life before assays. Mosquito larvae of *Aedes aegypti* (Gainesville) and *Culex pipiens* (Buckeye strain)^52^ were fed fish food (Tetramin, Melle, Germany), and adults were fed with water and 10% sucrose solution *ad libitum*. *Rhodnius prolixus* were obtained from BEI resources and fed rabbit blood (Pel-Freez Biologicals, Rogers, AZ, USA) in each instar through Hemotek artificial feeding membrane (Hemotek, Blackburn, UK). *Mezium affine* were obtained from the Ohio State University Insectary and fed oat meals *ad libitum*^53^. Fruit flies were obtained from Bloomington *Drosophila* Stock Center and fed with cornmeal food *ad libitum* (refer to Table S1 for the animals used in each figure).

### Experimental setup for humidity entrainment and free-running conditions

Of importance, we conducted preliminary studies using two incubator systems to control humidity cycles (Percival Scientific and Power Scientific). Vibration caused by the humidifiers and heating/cooling systems was sufficient to affect activity and behavior, reducing the level of entrainment to humidity. These issues required the development of a more robust, larger system to move the insects away from thermal and humidity control apparatuses (Fig. S3). To do so, the humidity entrainment is conducted in a greenhouse (6-ft L x 6-ft W x 6-ft H, FlowerHouse, Clio, MI). The cycles of low (40-50%RH, arid phase, A) and high (>85%RH, humid phase, H) humidity levels for entrainment are 12h A:12h H under constant light (350 lux) and temperature (25°C) conditions unless otherwise stated. The system was placed in a small room that was only accessible through another room that only served as a buffer space to prevent any unknown factors from impacting insect behavior. The humidity levels (AH) were achieved through humidifiers (Honeywell, Charlotte, NC) and dehumidifiers (vremi, China) controlled by humidity controllers (Inkbird IHC200, Lerway Tech, China). In free-running conditions, constant high humidity (HH) was used to avoid the impact of dehydration stress on animals ^52^. The relative humidity, air pressure, and light intensity were monitored by a HOBO Analog/temperature/RH/Light Logger (MX1104, Onset, Bourne, MA) and a digital barometer (ThermoPro TP67-B Wireless Weather Station, ThermoPro, Lawrenceville, GA). All activity monitors were placed on independent shelf units with vibration pads beneath each to prevent minor vibrations. Air venting from the humidifiers and dehumidifiers was positioned to avoid direct contact with the activity monitors. With this system, light, temperature, and barometric pressure remained constant, but humidity varied by 40-50% upon experimental needs (Fig. 1 and Supplementary Fig. 1).

### Locomotor activity and sleep assessment

The locomotor activities of insects in the Locomotor Activity Monitoring System (LAM25) were simultaneously measured and recorded with the DAMSystem3 Data Collection Software (TriKinetics, Waltham, MA). The actograms were generated and plotted by ImageJ plug-in ActogramJ^54^. General locomotor activity and sleep from two days before and after humidity entrainment were analyzed using the rethomics platform in R with behavr, damr, scopr, ggetho, sleepr, and zeitgeber packages as previously described ^26,55,56^. The statistics used for the comparison of the 24-h fly activities (Fig. 2B) and insect activities in arid and humid phases (Fig. 2C) were one-way ANOVA with post-hoc Tukey HSD test and Student’s t-test, respectively. As the periodogram analysis of the first three days of the free-running conditions, we first used the Lomb-Scargle periodogram in ActogramJ to determine the rhythmicity if a peak was above the 0.05 significance line and then the χ2 periodogram to calculate the period length and power^57^. The Fisher’s exact test and post hoc Bonferroni test were used to assess statistical significance when comparing the rhythmicity between wild types and mutants. Sleep was quantified as inactivity longer than five minutes using Rethomics and the aforementioned R packages as previously described ^55,58,59^.

## Acknowledgments

We thank the staff at the University of Cincinnati for their assistance in developing the mesocosm and devices used in this research.

## Funding

The research reported in this publication was supported by the National Institute of Allergy and Infectious Diseases under Award Numbers R01AI148551, R21AI166633, and R21AI176098 (JBB). The content is solely the responsibility of the authors and does not necessarily represent the official views of the National Institutes of Health.

## Author contributions

Conceptualization, Writing – review & editing, Visualization, Data curation, Investigation, Supervision, Formal analysis: S.C.C. and J.B.B. Funding acquisition: J.B.B. Investigation: G.C., D.E., N.C., T.C., H.T., L.E.L., L.W., E.G., J.S., J.P., L.H., A.B., E.T., J.T., N.G., A.Y. Formal analysis: E.G., J.S., J.P., L.H., A.B., E.T., J.T., N.G.

## Competing interests

The authors declare that they have no competing interests.

## Data and materials availability

Data will be made available on Dryad at acceptance

## Notes

### Competing Interest Statement

The authors have declared no competing interest.

### Summary of Updates

We have added two additional experiments.

