## Supplemental materials for "Humidity as a potential zeitgeber for circadian entrainment of insect systems"

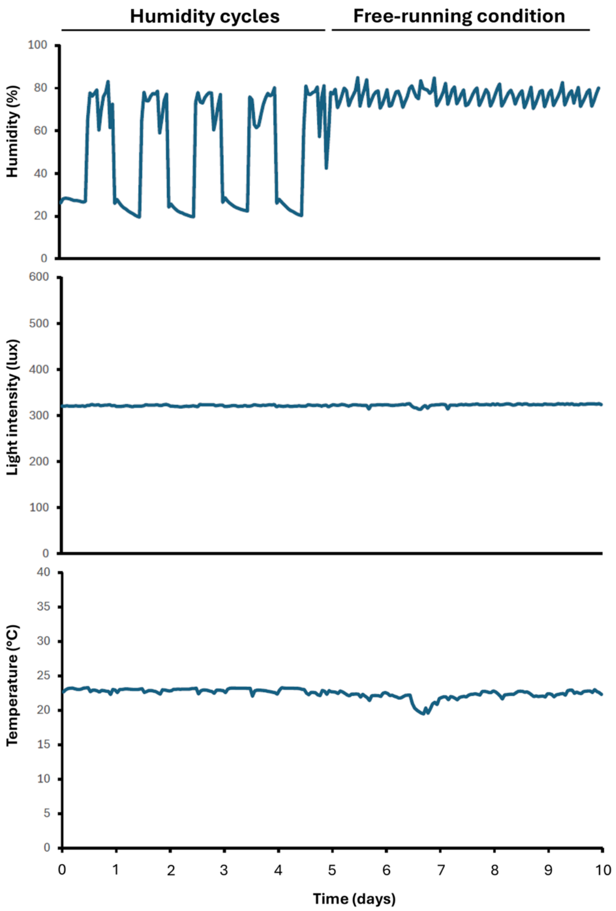


**Supplementary Fig. 1.** One representative recording shows humidity (top), light intensity (middle), and temperature (bottom) during the humidity cycles (12h: 12h A:H) and under free-running conditions. Barometric pressure was checked daily and remained between 780 and 785 mm Hg.

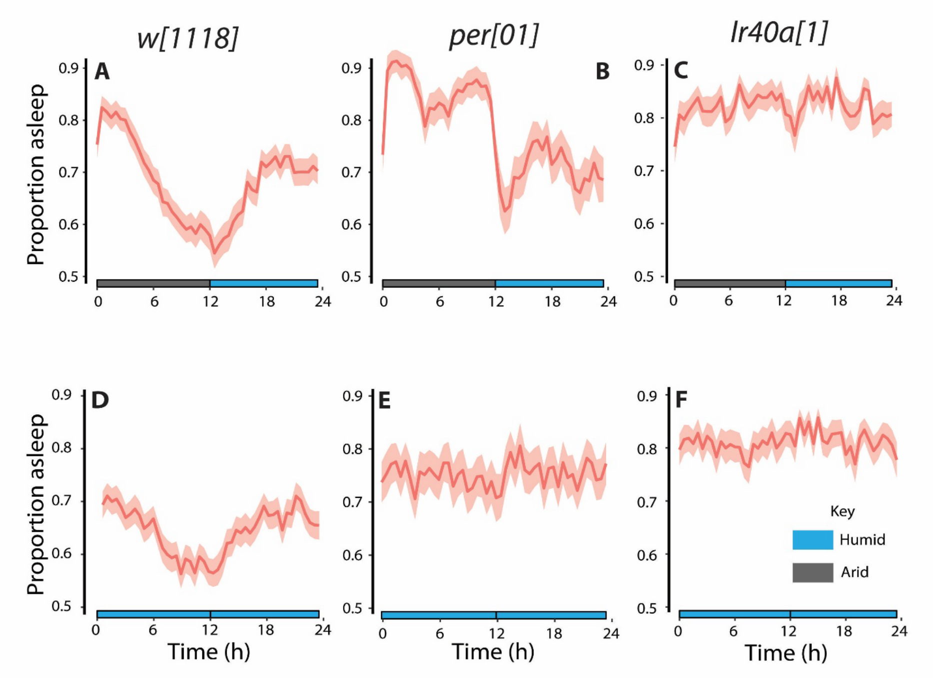


**Supplementary Fig. 2.** Sleep patterns of *Drosophila.* Wild type (*w*[1118], ♂, n=44) and mutants (*per*[01], ♂, n=46; *Ir40a*[1], ♂, n=48) in humidity cycles (AH; A-C) and the free-running conditions (HH; D-F).

Commented [BJ1]: Move to supplement

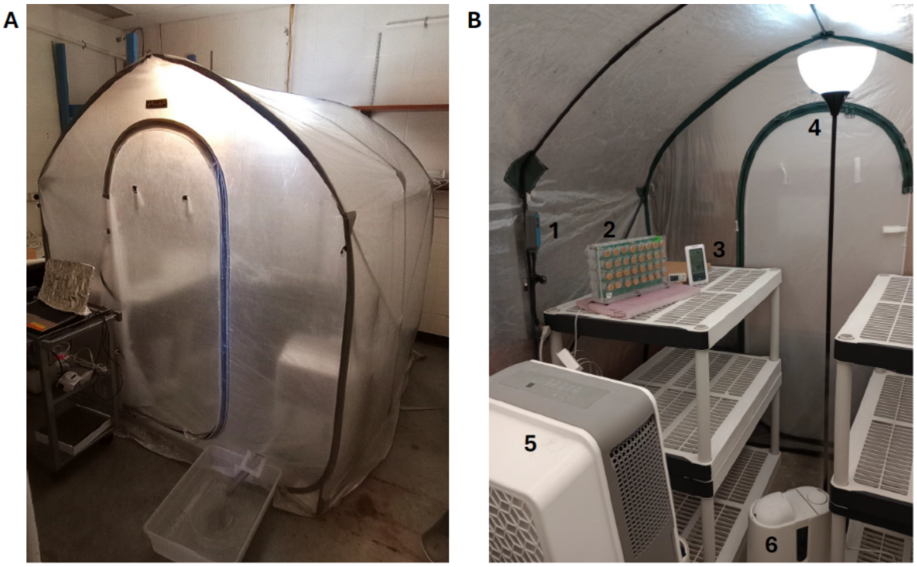


**Supplementary Fig. 3.** Experimental setup for humidity entrainment of insect circadian clocks. A: The humidity entrainment is carried on in a vibration-free greenhouse. B. The setup for humidity cycles. 1: humidity controller, 2: Locomotor Activity Monitoring System (LAM25), 3: monitors for light/temperature/humidity/air pressure, 4: lamp, 5: dehumidifier, and 6: humidifier.

Table S1 for the animals used in each figure.

| Figure | Species or designation | Source | Identifiers |
|---|---|---|---|
| Fig. 4 | *D. melanogaster* (Oregon-R-C, ♂) | Bloomington Drosophila Stock Center | RRID:BDSC_5 |
| | *D. melanogaster* ($w^{1118}$, ♂) | Bloomington Drosophila Stock Center | RRID:BDSC_5905 |
| | *D. melanogaster* (Canton S, ♂) | Bloomington Drosophila Stock Center | RRID:BDSC_64349 |
| | *R. prolixus* (♀) | BEI Resources | |
| | *M. affine* (sex unknown) | Benoit Lab | |
| | *Ae. aegypti* (♀) | BEI Resources | |
| | *C. pipiens* (♀) | Benoit Lab | Buckeye strain from Northern Ohio |
| Fig. 7 | $per^{01}$ (♂) | Bloomington Drosophila Stock Center | RRID:BDSC_80928 |
| | $tim^{01}$ (♂) | Bloomington Drosophila Stock Center | RRID:BDSC_80930 |
| | $cyc^{01}$ (♂) | Bloomington Drosophila Stock Center | RRID:BDSC_80929 |
| | $Clk^{Jrk}$ (♂) | Bloomington Drosophila Stock Center | RRID:BDSC_80927 |
| | $pdf^{01}$ (♂) | Bloomington Drosophila Stock Center | RRID:BDSC_26654 |
| Fig. 8 | $Ir40a^{1}$ (♂) | Bloomington Drosophila Stock Center | RRID:BDSC_81252 |
| | $Ir25a^{2}$ (♂) | Bloomington Drosophila Stock Center | RRID:BDSC_41737 |

| | | | |
|---|---|---|---|
| | *Df(Ir25a)* (♂) | Bloomington Drosophila Stock Center | RRID:BDSC_7496 |
| | $Ir93a^{MI0555}$ (♂) | Bloomington Drosophila Stock Center | RRID:BDSC_42090 |
| | $Tmem63^{2}$ (♂) | Bloomington Drosophila Stock Center | RRID:BDSC_92652 |
| | $Obp59a^{1}$ (♂) | Bloomington Drosophila Stock Center | RRID:BDSC_80683 |
| | $Ir68a^{MB05565}$ (♂) | Bloomington Drosophila Stock Center | RRID:BDSC_26031 |
| | $Ir76b^{1}$ (♂) | Bloomington Drosophila Stock Center | RRID:BDSC_51309 |
| | $ppk28^{G981}$ (♂) | Bloomington Drosophila Stock Center | RRID:BDSC_33559 |
| | $Orco^{2}$ (♂) | Bloomington Drosophila Stock Center | RRID:BDSC_23130 |
| Fig. 4 | $w^{1118}$ (♂) | Bloomington Drosophila Stock Center | RRID:BDSC_5905 |
